# High-Speed Super-Resolution Imaging Using Protein-Assisted DNA-PAINT

**DOI:** 10.1101/2020.02.11.943506

**Authors:** Mike Filius, Tao Ju Cui, Adithya Ananth, Margreet Docter, Jorrit W. Hegge, John van der Oost, Chirlmin Joo

## Abstract

Super-resolution imaging allows for visualization of cellular structures on a nanoscale level. DNA-PAINT (DNA Point Accumulation In Nanoscale Topology) is a super-resolution method that depends on the binding and unbinding of DNA imager strands. The current DNA-PAINT technique suffers from slow acquisition due to the low binding rate of the imager strands. Here we report on a method where imager strands are loaded into a protein, Argonaute (Ago), that allows for faster binding. Ago pre-orders the DNA imager strand into a helical conformation, allowing for 10 times faster target binding. Using a 2D DNA origami structure, we demonstrate that Ago-assisted DNA-PAINT (Ago-PAINT) can speed up the current DNA-PAINT technique by an order of magnitude while maintaining the high spatial resolution. We envision this tool to be useful not only for super-resolution imaging, but also for other techniques that rely on nucleic-acid interactions.

---

Single-molecule localization microscopy techniques allow researchers to image cellular structures that are not visible through diffraction-limited microscopy methods. Most single-molecule localization techniques rely on the stochastic blinking of fluorescent signal, by using photoswitchable fluorophores as in photoactivated-localization microscopy (PALM)^1^ and (direct) stochastic optical reconstruction microscopy ((d)STORM)^2^. An alternative approach to achieve stochastic blinking is through fluorescent probes that transiently bind their target, as in point accumulation in nanoscale topography (PAINT).^3–5^

In DNA-PAINT, a fluorophore is attached to a short DNA oligonucleotide (or imager strand) that specifically binds to a complementary target DNA sequence (or docking strand).^6^ The stochastic blinking of signals is achieved through binding and unbinding of the incoming imager strands to the docking strands and is imaged using total internal reflection fluorescence (TIRF). By changing the length and sequence of an imager strand, one can tune the on- and off-rates of the imager and adjust the specificity. This allows for high multiplexing capabilities since the number of probes is only limited by the number of orthogonal DNA sequences. Furthermore, compared to conventional super-resolution techniques, DNA-PAINT comes with the advantage that imager strands are continuously replenished from the solution and thus photobleaching is circumvented during the imaging process.

A critical limitation of DNA-PAINT, however, is the low binding rate of DNA, which is typically in the order of 10^6^ M^−1^ s^−1^. Given this binding rate, obtaining images with high spatial resolution (5 nm) usually takes several hours.^7–9^ Shorter acquisition times can be achieved by increasing concentration of the imager strand.

However, single-molecule binding events become unresolvable from the background of unbound imager strands, even when using TIRF. To reduce this acquisition time, DNA-PAINT has recently been combined with single-molecule Förster Resonance Energy Transfer (smFRET).^10,11^ This, however, comes at a cost of reduced spatial resolution due to limited energy transfer efficiency between donor and acceptor dyes and camera sensitivity being a limiting factor for dyes used in the far red spectrum. Here we describe an alternative approach, in which protein-assisted delivery of imager strands is demonstrated to speed up the acquisition time 10-fold and only requires a single fluorescent channel.

Argonaute proteins (Agos) are a class of enzymes that utilize a DNA or RNA guide to find a complementary target, either to inactivate or to cleave it. In eukaryotes, an RNA guide directs Ago to complementary RNA targets for post-transcriptional regulation.^12^ Ago proteins initially bind their target through base pairing with the seed segment of the guide (nucleotides 2-7 for human Ago).^13–15^ Crystal structures have revealed that Ago pre-orders this seed segment into a helical conformation, allowing for the formation of a double helix between guide and target, and hence effectively pre-paying the entropic cost of target binding.^16,17^ This results in binding rates that are near-diffusion limited (~10^7^ M^−^ 1 s^−1^).^18–21^ In prokaryotes, there is a broad diversity of Agos with respect to the identity of their guide (RNA/DNA) and their target (RNA/DNA).^22,23^ Some well-characterized prokaryotic Ago nucleases (*Thermus thermophilus* Ago and *Clostridium butyricum* Ago) use DNA guides to target single-stranded (ss)DNA.^24,25^

Here we describe a new DNA-PAINT method based on protein-assisted delivery of DNA imager strands, which allows for faster acquisition of super-resolved nanostructures. In this Ago-PAINT method, we use a wildtype Ago protein from the bacterium *Clostridium butyricum* (*Cb*Ago) to speed up the kinetic binding of DNA imager strands. *Cb*Ago reshapes the binding landscape of the imager strand, resulting in a 10-fold higher binding rate compared to conventional DNA-PAINT. In addition, we show that one can implement Ago-PAINT with minimal imager strand complexity whilst retaining the programmability and specificity of DNA-PAINT, due to the favourable targeting feature of *Cb*Ago.^25,26^ We determine the spatial resolution of Ago-PAINT through the use of 2D DNA origami structures and show that Ago-PAINT generates super-resolution images of diffraction limited structures at least 10-fold faster than conventional DNA-PAINT.

## RESULTS

For high-quality super-resolution images, a PAINT-based method requires more than five transient binding events per localization spot^7^, each with a dwell time of at least several hundreds of milliseconds.^7–9^ A typical 8-nt DNA-PAINT imager strand exhibits an on-rate (*k*_*on*_) of ~ 10^6^ M^−1^ s^−1^ and a dwell time (an inverse of an off-rate, 1/*k*_*off*_) of ~1 s.^9^ DNA-PAINT experiments use an imager strand concentration between 1-10 nM. This range is chosen to be high enough to obtain a sufficient number of binding events during the acquisition time, but not too high to avoid cross-talk localization between structures.^7^

We determined the on- and off-rates of Ago-PAINT imager strands and compared these to the on- and off-rate of conventional DNA-PAINT with the same imager strands using a smFRET assay (**Figure 1)**. Acceptor (Cy5)-labelled ssDNA targets were immobilized through biotin-streptavidin conjugation on a PEGylated quartz slide. Next, either donor (Cy3)-labelled 8-nt DNA-PAINT imager strands or Ago-PAINT imager strands (*Cb*Ago loaded with a Cy3-labelled guide) were injected, and their interactions with the immobilized target strand were probed using TIRF microscopy (**Figure 1A**). The assay was designed to give a high-FRET signal upon specific binding of either DNA imager strand or Ago-guide complex to the complementary target (**Figure 1B and C**). The Cy3 position was chosen the same as in our previous studies on *Cb*Ago, where we showed the dye position does not bring any photophysical artefacts.^26,27^ The time between introduction of the imager strands and the first binding event is the arrival time (which is the inverse of the on-rate, *k*_*on*_). The duration of the FRET binding events is the dwell time (**Figure 1B**).

**Figure 1.**
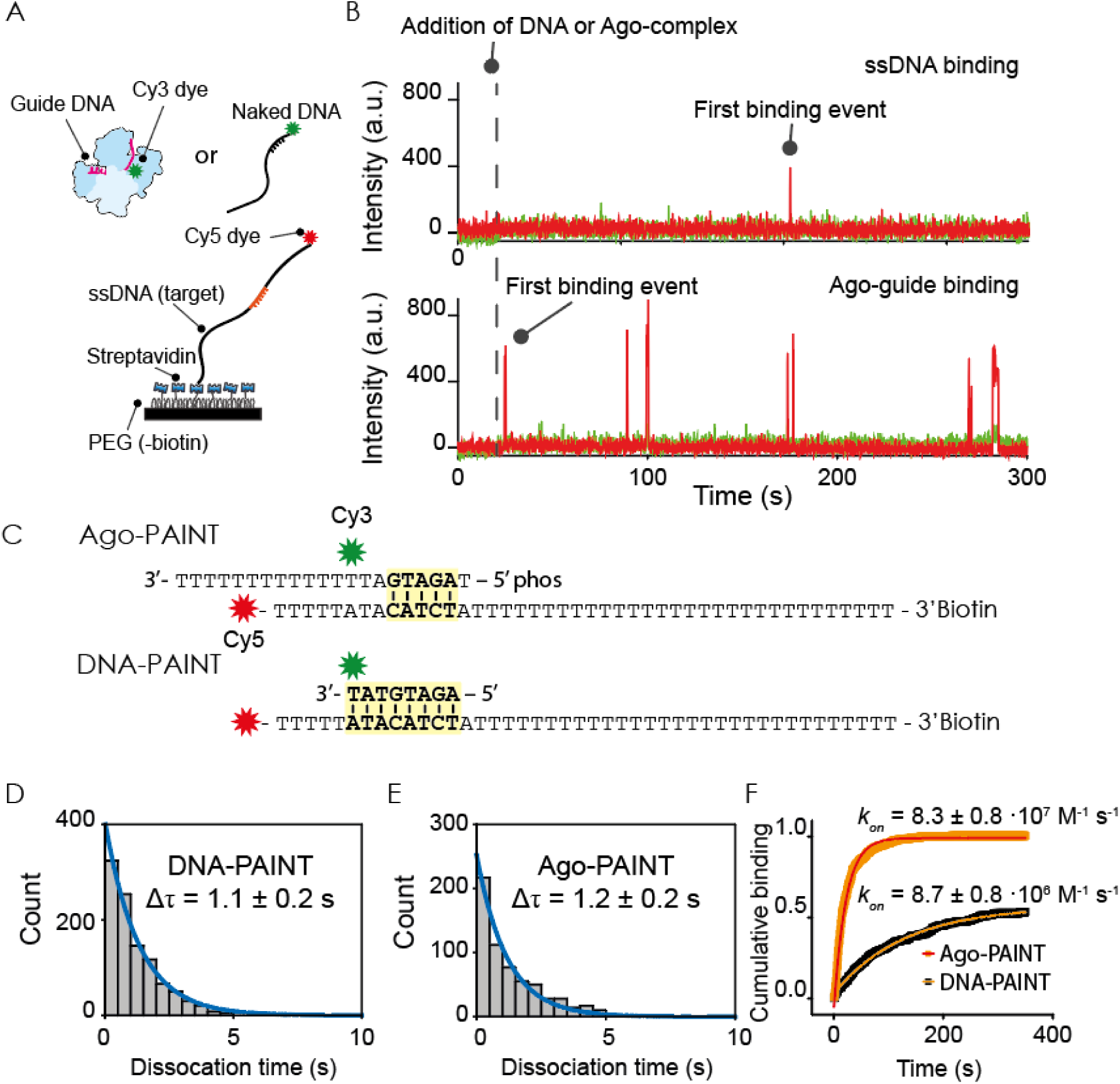
Single-molecule FRET assay to quantify binding kinetics Ago vs DNA-PAINT. (A) A schematic of the single-molecule FRET assay with the target strand immobilized on a PEGylated surface through biotin-streptavidin conjugation. The green and red stars indicate the Cy3 and Cy5 dye respectively. Binding of Ago-guide complex or ssDNA probe to the ssDNA target results in high FRET signal. (B) Representative traces of ssDNA binding (top) and Ago-complex binding (bottom). The dashed line indicates the timepoint at which Ago-guide or DNA is introduced inside the microfluidic chamber. (C) A schematic of the sequences used for Ago-PAINT and DNA-PAINT. Upon binding, both constructs will give rise to a high FRET signal. (D) Dwell-time histogram (Δτ) of ssDNA (sequence shown in Figure 1C). Maximum likelihood estimation (MLE) gives 1.1 ± 0.2 s as the parameter for a single-exponential distribution (blue line). Number of data-points: 1029. (E) Dwell-time histogram (delta T) of Ago (sequence shown in Figure 1A). MLE fitting gives 1.2 ± 0.2 s as the parameter for a single-exponential distribution (blue line). Number of data-points: 696. (F) Cumulative binding event plots of DNA-PAINT (Black) and Ago-PAINT (Orange) vs time. A single-exponential fit is used for DNA-PAINT (red line) and Ago-PAINT (orange line). Errors in (D), (E) and (F) are determined by taking the 95% confidence interval of 10^5^ bootstraps.

For a comparison between Ago-PAINT and DNA-PAINT, we designed an 8-nt DNA-PAINT imager strand (**Figure 1C**) and found that under our experimental conditions the average dwell time of this imager strand is 1.1 ± 0.2 s (**Figure 1D**). Next, we sought to find an Ago-PAINT guide with a similar dwell time. The first nucleotide of an Ago guide is embedded within the protein structure (**Figure S1A**).^16,17^ Therefore, we determined the dwell time of Ago-guide complexes with different numbers (N) of base pairing with the target starting from the second nucleotide onwards (**Figure S1B**). A guide with N=5 (nt 2-6) base-pairing to the target exhibited a comparable dwell time of 1.2 ± 0.2 s (**Figure 1E.** We observed that for Ago-PAINT the apparent binding rate is influenced by the number of base pairs that are formed between the guide and its target. For N=5 or larger, the on-rate reaches a saturated value (*k*_*on*_ = 0.6-1.0 · 10^8^ M^−1^ s^−1^) (**Figure S1C**). Those values are 10 times higher than the typical on-rates for an 8-nt DNA-PAINT imager strand, 8.7 ± 0.8 · 10^6^ M^−1^ s^−1^ (**Figure 1F**).

To demonstrate the use of Ago-PAINT for super-resolution imaging, we designed a rectangular 2-dimensional DNA origami structure of 76 nm x 80 nm (**Figure 2 and Figure S2**). The DNA origami structure has four docking sites that are spaced 61 nm x 68 nm apart (**Figure 2A**). To achieve optimal Ago binding to the DNA origami docking strands, we introduced a polyT linker (T_30_) between the target sequence of Ago and the DNA origami structure (**Figure 2A**, right panel).

**Figure 2.**
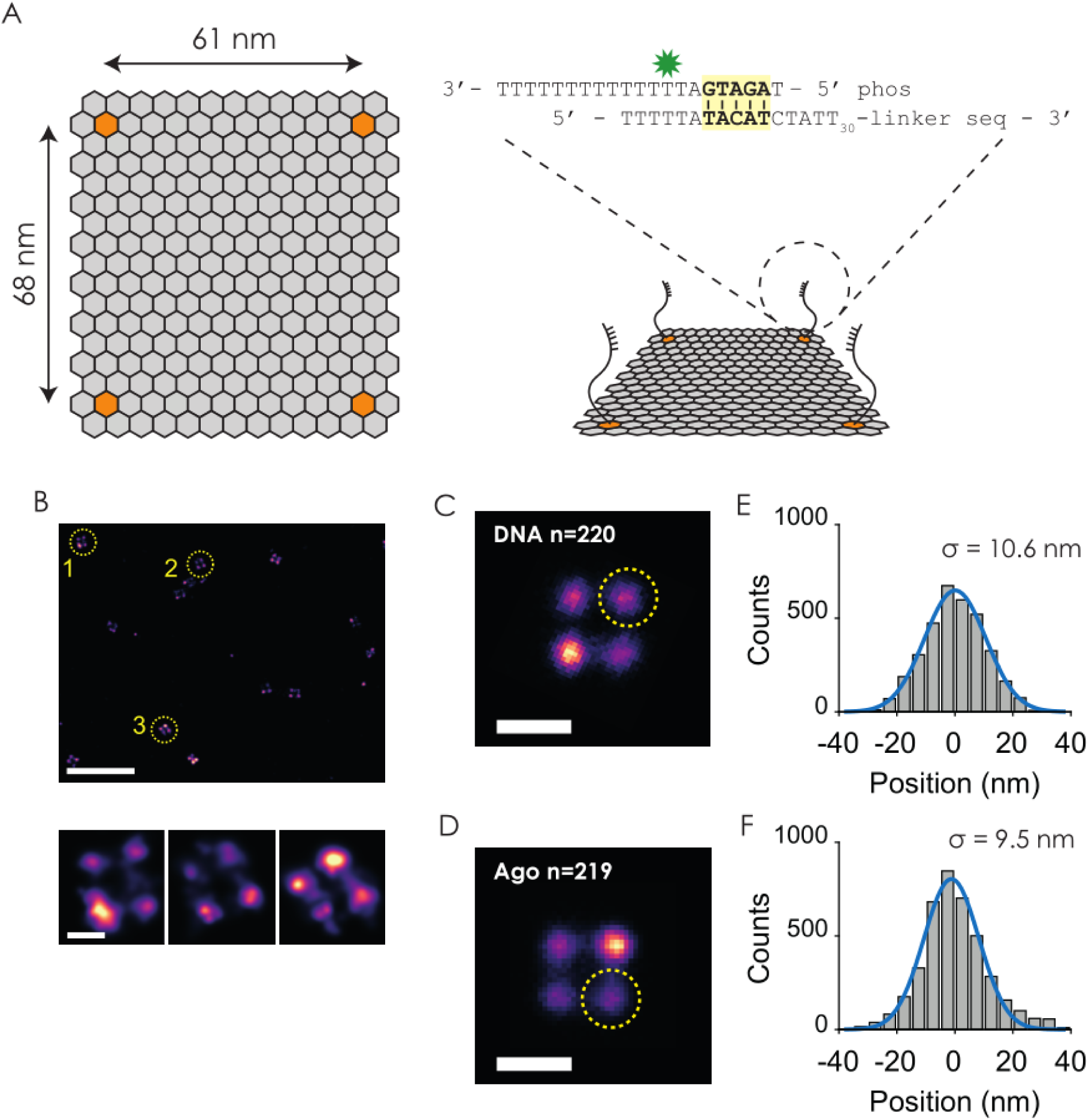
Ago-PAINT enables the same localization precision as conventional DNA-PAINT. (A) Left: A schematic design of the 2D-DNA origami structure. The orange honeycombs indicate the approximate locations of binding sites. Right: 3D representation of the imaging scheme with the used docking strand sequence. The green star indicates the position of the Cy3 dye labelled on the backbone of an amino-modified thymine. (B) A representative super-resolution image showcasing binding sites of the 2D-DNA origami structures using Ago-PAINT. Bottom: Super-resolution reconstruction of the four-corner origami structures of the top panel. (C) A summed image of 220 origami structures visualized through the use of DNA-PAINT. (D) A summed image of 219 origami structures made through the use of Ago-PAINT. The concentration of imager strand was 1 nM for both conventional DNA-PAINT and Ago-PAINT. (E) Fitting of a cross-sectional intensity histogram from the yellow encircled area in Figure (C) to a Gaussian (blue line) shows that a localisation precision of 10.6 nm can be achieved, similar to Ago-PAINT. (F) Fitting of a cross-sectional intensity histogram from the yellow encircled area in Figure (D) to a Gaussian (blue line) shows that a localisation precision of 9.5 nm is possible under these conditions. Scale bars in (B) indicate 500 nm (top) and 50 nm (bottom three). Scale bars in (C) and (E) indicate 100 nm.

Next, we sought to compare the localization precision of Ago-PAINT and DNA-PAINT. We tested our Ago-PAINT approach by injecting guide-loaded Ago into our flow cell in which DNA origami structure were immobilized. A super-resolution image could be reconstructed from the Ago-PAINT data which revealed four detectable spots on the origami structures as expected from our assay design (**Figure 2B**). We determined the localization precision by selecting 220 origami structures for DNA-PAINT and 219 structures for Ago-PAINT and created a sum image using the Picasso analysis software^7^ (**Figure 2C and 2D**). The localization precision was determined by plotting the cross-sectional histogram of one of the four binding sites of the summed DNA origami structure. For DNA-PAINT this resulted in a localization precision of 10.6 nm (**Figure 2E**) and for Ago-PAINT we found a localization precision of 9.5 nm (**Figure 2F**). The histogram demonstrates that Ago-PAINT delivers the same quality of localization precision when compared to the DNA-PAINT approach. Analysis of the data based on nearest neighbour analysis^28^ reconfirms this finding since a localization precision of 9.7 nm was found for both Ago-PAINT and DNA-PAINT (**Figure S3**).

Furthermore, we determined the possibility to use different linker lengths for Ago-PAINT imaging. With this in mind, we designed DNA origami structures with longer linkers (50 thymines or 100 thymine nucleotides) and found that this did not affect the localization precision of Ago-PAINT (**Figure S4 and S5**), showing that Ago-PAINT is compatible with various linker lengths (≥ T30). When we used a shorter linker (5 thymines), we could not register a sufficient number of binding events (data not shown).

Finally, we compared the speed of super-resolution imaging through Ago-PAINT with the conventional DNA-PAINT approach using the 2D DNA origami structures as a testing platform. We evaluated the quality of a super-resolution image after each time-point for both Ago-PAINT and DNA-PAINT (**Figure 3A**). The overall resolution of a single-molecule localization microscopy (SMLM) image is dependent on the number of localizations (binding events) per docking strand 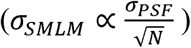. Therefore, to quantify the speed of imaging we plot the standard error of the localization precision as a function of frame number (**Figure 3B**) where we took the sigma values from **Figure 2E and 2F** as the localization precision. We observed that the standard error of the localization precision for Ago-PAINT is smaller than that of DNA-PAINT at each time point, indicating that super-resolved images of identical resolution will be obtained 10 times faster through Ago-PAINT compared to DNA-PAINT. This result is further supported by the intensity vs time traces, which shows that our Ago-PAINT method results in more binding events compared to DNA-PAINT approach, under similar conditions with DNA concentrations of 1 nM (**Figure 3C-E and Figure S6**). The on-rates for both Ago-PAINT (*k*_*on*_ = 4.4 ± 0.1 ·10^7^ M^−1^ s^−1^) and DNA-PAINT (*k*_*on*_ = 6.6 ± 0.1 · 10^6^ M^−1^ s^−1^) on our DNA-origami structure **(Figure 3E)** are similar to the on-rates that we found in our single-molecule experiments (**Figure 1F**).

**Figure 3.**
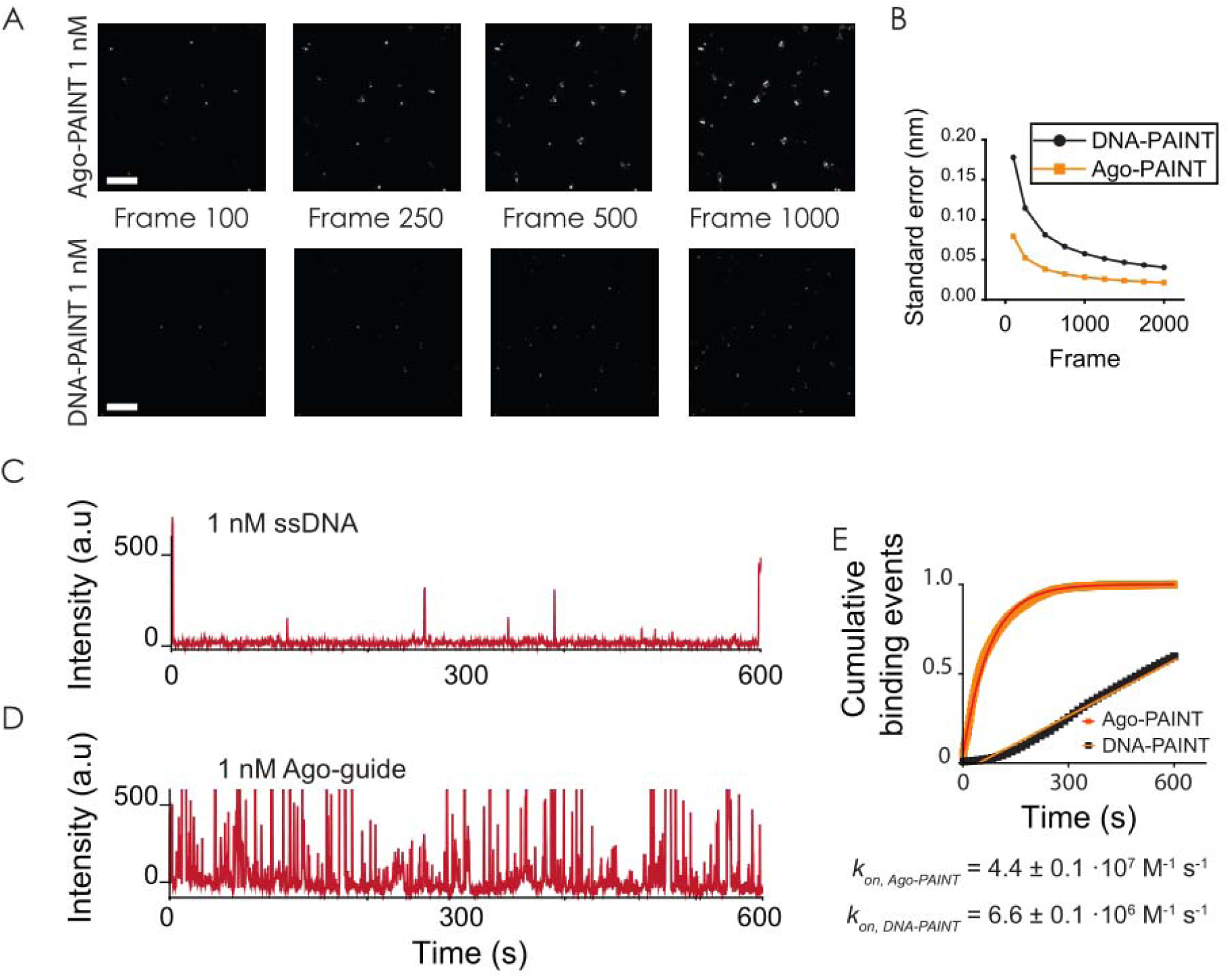
Ago-PAINT enables fast imaging of super-resolved structures. (A) Snapshots in time for Ago-PAINT (top) and DNA-PAINT (bottom) showing super-resolution images being formed over time. Exposure time: 0.3 s. The same color scale is used for the intensity in all images. (B) Standard error of Ago-PAINT vs DNA-PAINT plotted versus frame number. (C) Representative intensity vs time data trace of DNA-PAINT at 1 nM DNA concentration shows few binding events occurring within 600 s. The raw data trace is taken from a single origami plate. (D) Representative intensity vs time data trace of 1 nM Ago-guide complex shows binding events occurring frequently within 600 s. The raw data trace is taken from a single origami plate. (D) Normalized cumulative distribution of dark times (the time between binding events) for DNA-PAINT (black, n = 4870) and Ago-PAINT (orange, n = 5793). A single-exponential growth curve (red for DNA-PAINT, orange for Ago-PAINT) is used to estimate the binding rate. Scale bars in (A) indicate 500 nm.

## DISCUSSION

Here we presented a proof-of-concept of Ago-PAINT that allows for rapid super-resolution imaging. We demonstrated that fast Ago-PAINT recording can be used to acquire super-resolution images of nanostructures while retaining the programmability and predictability of DNA-PAINT.

In order to fully visualize real-time interactions between multibody cellular components, one would ideally want to look at multiple components at the same time. Effort has been put into temporal^29^ or spectral^30^ multiplexing of DNA-PAINT technology. In our previous work, we showed that different guide sequences resulted in distinctly different binding kinetics.^25^ These kinetic fingerprints will allow for additional freedom when designing Ago-PAINT.^29,31^ Furthermore, optimization of the imager sequence and imaging conditions allowed for an order of magnitude faster imaging for conventional DNA-PAINT^32^. We expect that optimization of the guide sequence and buffer conditions could further improve the kinetics Ago-PAINT. In this study, Ago-PAINT experiments are performed with the wild-type *Cb*Ago protein which substantially increases the probe size compared to conventional DNA-PAINT. However, successful applications of Argonaute proteins for *in vivo* gene silencing^33,34^ hint that our Ago-PAINT approach could be used in cellular super-resolution imaging. While targeting complex cellular structures in cells could be an issue with a full size *Cb*Ago, it is possible to use truncated versions of Ago. Some truncated versions of approximately half the size (short Agos) exist in nature^22^. We speculate that it will be possible to truncate them further as Ago-PAINT only relies on the property of pre-forming the helix structure of the imager strand and a variant from *Kluyveromyces polysporus* that contains only the C-lobe was reported to retain almost all the binding properties of the untruncated version.^35^ Furthermore, as the imager strand is loaded and protected inside the protein, degradation of the imager strand is less likely to occur over time, unlike oligos that are rapidly digested.^36^

In this paper we demonstrated the use of *Cb*Ago for super-resolution microscopy. While this *Cb*Ago targets ssDNA, Agos from other species can target RNA^22^. For example, the Ago from *Marinitoga piezophila* (*Mp*Ago)^37,38^ targets RNA and one could harness the property of a high association rate for other single molecule imaging applications such as RNA sensing. Recently, *dTt*Ago has been combined with FISH^39^ to allow for labelling of genomic loci in fixed cells. We anticipate the use of RNA guided Agos for a significant speed-up in similar applications for RNA FISH. Lastly, complementary approaches such as DNA-based STED imaging^40^, FRET-PAINT^30^, qPAINT^41^ or crosslinking on single-molecule target using Action-PAINT^42^ could be combined with our Ago-PAINT approach. We envision the use of Ago-PAINT as a general toolkit to speed up many current existing applications that rely on base-pairing interactions.

## MATERIALS AND METHODS

### Expression and purification of *Cb*Ago

The *Cb*Ago gene was codon harmonized for E.coli Bl21(DE3) and inserted into a pET-His6 MBP TEV cloning vector (Addgene plasmid #29656) using ligation independent cloning. The *Cb*Ago protein was expressed in E.coli Bl21(DE3) Rosetta™ 2 (Novagen). Cultures were grown at 37 °C in LB medium containing 50 μg ml^−1^ kanamycin and 34 μg ml^−1^ chloramphenicol till an OD600 nm of 0.7 was reached. *Cb*Ago expression was induced by addition of isopropyl β-D-1-thiogalactopyranoside (IPTG) to a final concentration of 0.1 mM. During the expression cells were incubated at 18 °C for 16 h with continues shaking. Cells were harvested by centrifugation and lysed, through sonication (Bandelin, Sonopuls. 30% power, 1 s on/2 s off for 5 min) in lysis buffer containing 20 mM Tris-HCl pH 7.5, 250 mM NaCl, 5 mM imidazole, supplemented with a EDTA free protease inhibitor cocktail tablet (Roche). The soluble fraction of the lysate was loaded on a nickel column (HisTrap Hp, GE healthcare). The column was extensively washed with wash buffer containing 20 mM Tris-HCl pH 7.5, 250mM NaCl and 30 mM imidazole. Bound protein was eluted by increasing the concentration of imidazole in the wash buffer to 250 mM. The eluted protein was dialysed at 4 °C overnight against 20 mM HEPES pH 7.5, 250 mM KCl, and 1 mM dithiothreitol (DTT) in the presence of 1 mg TEV protease (expressed and purified according to Tropea et al.^43^) to cleave of the His6-MBP tag. Next the cleaved protein was diluted in 20 mM HEPES pH 7.5 to lower the final salt concentration to 125 mM KCl. The diluted protein was applied to a heparin column (HiTrapHeparin HP, GE Healthcare), washed with 20 mM HEPES pH 7.5, 125 mM KCland eluted with a linear gradient of 0.125–2 M KCl. Next, the eluted protein was loaded onto a size exclusion column (Superdex 200 16/600 column, GE Healthcare) and eluted with 20 mM HEPES pH 7.5, 500 mM KCl and 1 mM DTT.

Purified *Cb*Ago protein was diluted in size exclusion buffer to a final concentration of 5 μM. Aliquots were flash frozen in liquid nitrogen and stored at −80 °C.

### Single-molecule setup

All experiments were performed on a custom-built microscope setup. An inverted microscope (IX73, Olympus) with prism-based total internal reflection is used. In combination with a 532 nm diode laser (Compass 215M/50mW, Coherent). A 60x water immersion objective (UPLSAPO60XW, Olympus) was used for the collection of photons from the Cy3 and Cy5 dyes on the surface, after which a 532 nm long pass filter (LDP01-532RU-25, Semrock) blocks the excitation light. A dichroic mirror (635 dcxr, Chroma) separates the fluorescence signal which is then projected onto an EM-CCD camera (iXon Ultra, DU-897U-CS0-#BV, Andor Technology). A series of EM-CDD images was recorded using custom-made program in Visual C++ (Microsoft). Time traces were extracted from the EM-CDD images using IDL (ITT Visual Information Solution) and further analyzed with Matlab (Mathworks) and Origin (Origin Lab).

### Single-molecule data acquisition

To avoid non-specific binding of *Cb*Ago protein to the surface, quartz slides were PEGylated as previously described (Chandradoss 2014). Briefly, acidic piranha etched quartz slides (Finkenbeiner) were passivated twice with polyethylene glycol (PEG). The first round PEGylation was performed with mPEG-SVA (Laysan) and PEG-biotin (Laysan), followed by a second round of PEGylation with MS(PEG)_4_ (ThermoFisher). After assembly of a microfluidic chamber, the slides were incubated with 1 % Tween-20 for 15 minutes. Excess Tween-20 was washed away with 100 µL T50 (50mM Tris-HCl, pH 8.0, 50 mM NaCl) followed by a 2 min incubation of 20 µL streptavidin (0.1 mg/mL, ThermoFisher). Excess streptavidin was removed with 100 µL T50. Next, for single-molecule experiments we immobilized 50 µL of 100 pM Cy5 labelled target DNA for 2 minutes, unbound DNA was washed with 100 µL T50, followed by 100 µL of origami-buffer (50 mM Tris-HCl, pH 8.0, 50 mM NaCl, 1 mM MnCl_2_, 5 mM MgCl_2_). The Ago-guide complex was formed by incubating 10 nM *Cb*Ago with 1 nM of Cy3 labelled DNA guide for 20 minutes at 37 °C in the origami-buffer. For single-molecule experiments, we injected 50 µL of 1 nM Ago-guide complex or 50 µL of 1 nM DNA-PAINT imager strand in imaging buffer (50 mM Tris-HCl, pH 8.0, 50 mM NaCl, 1 mM MnCl_2_, 5 mM MgCl_2_, 0.8 % glucose, 0.5 mg/mL glucose oxidase (Sigma), 85 ug/mL catalase (Merck) and 1 mM Trolox (Sigma)). The single-molecule FRET experiments for **Figure 1** were performed at room temperature (23 ± 2 °C). For super-resolution DNA origami experiments, we flushed 50 µL of ~200 pM DNA origami structures in a streptavidin coated channel and incubated for 3 minutes to allow for specific immobilization.

Unbound DNA-origami was washed with origami-buffer. Next, 50 µL of 100 pM of Ago-guide complex or 1 nM DNA-PAINT imager strand was injected in imaging buffer.

### Assembly of DNA oligo plate

The 2D rectangular DNA origami structure was designed by using CaDNAno software based on square lattice.^44^ The 2D rectangular DNA origami structure was twist corrected and structural behaviour of the origami plate was checked by coarse-grained simulations in CanDo.^45,46^ The parameters used for simulations are axial rise per base-pair = 0.34 nm, helix diameter = 2.25 nm, crossover spacing = 10.5 bp, axial stiffness = 1100 pN, bending stiffness = 230 pNnm^2^, torsional stiffness = 460 pN nm^2^, nick stiffness factor = 0.01. The 2D rectangular DNA origami structure self-assembled in a total reaction volume of 100 µL containing 10 nM of p8064 scaffold strand (Tilibit nanosystems), 100 nM core staples (Integrated DNA Technologies), 100 nM Ago-PAINT handles and 100 nM biotin handles in 1x TE folding buffer (Tilibit nanosystems) supplemented with 11 mM MgCl_2_. The origami structures were annealed using a thermocycler. First, the reaction mixture was heated for 10 minutes at 65 °C, then a temperature gradient was applied from 60 °C to 40 °C with a rate of 1 °C/hour. After self-assembly, the origami structures were purified using Amicon spin filter (100K MWCO) and stored in T50 buffer containing 11 mM MgCl_2_. The purified DNA origami structures were analysed on a 2 % agarose gel (Tris-borate-EDTA, 11 mM MgCl_2_). The gel was run at 90 V for 2 hours in ice. After staining the gel with ethidium bromide, the samples were imaged to verify the quality of the folding procedure (**Figure S2A**). Next, the purified origami sample were checked for rectangular structure by atomic force microscopy (AFM) on mica surface according to AFM imaging procedures. Briefly, 0.01 % (w/v) polylysine was incubated 1 min on a freshly cleaved 3 mm (1/8 inch) diameter mica disk. The mica surface was gently washed with MQ water and blow dried with N_2_. Next, 5 µl of 500 pM DNA origami samples was incubated onto a mica disk for 5 minutes. The mica disk was washed gently with 1 ml (3x) of folding buffer with 11 mM MgCl2 to remove any unbound DNA origami structures, then quickly rinsed with MQ water and blow dried with N2. Dry AFM images were acquired in Bruker Multimode 8 AFM. Sharp AFM tips were used for AFM measurements (Bruker PeakForce HIRS-F-B) with 0.12 N/m nominal spring constant. AFM images were acquired in tapping mode. Example images of AFM images can be found in **Figure S2B and C**.

### Super-resolution data analysis

CCD movies were acquired through custom-written program. The resulting files were converted to .raw file format using a custom-written script in Matlab (Mathworks). Super-resolution reconstruction, drift-correction and alignment were performed using the Picasso software package,^7^ for both Ago-PAINT and DNA-PAINT.

## CONTRIBUTIONS

M.F., T.J.C., C.J. conceived the project and designed the experiments. M.F., T.J.C., A.A. and C.J. designed and made the 2D DNA origami structure. J.W.H. and J.v.d.O. expressed and purified *Cb*Ago. M.F. and T.J.C. performed all single-molecule and DNA origami experiments. M.F., T.J.C., M.D. analysed the data. M.F., T.J.C. and C.J. wrote the manuscript. All authors read and improved the manuscript.

## ACKNOWLEDGEMENTS

We thank Carolien Bastiaanssen and Sung Hyun Kim for critical reading and feedback. C.J. was supported by Vidi (864.14.002) of the Netherlands Organization for Scientific Research; an ERC Consolidator Grant (819299) of the European Research Council; and Vrije Programma (SMPS) of the Foundation for Fundamental Research on Matter

